# Rapid shift in microbial community structure in a neutral hydrothermal hot spring from Costa Rica

**DOI:** 10.1101/2020.11.23.395137

**Authors:** Diego Rojas-Gätjens, Alejandro Arce-Rodríguez, Fernando Puente-Sánchez, Roberto Avendaño, Eduardo Libby, Geraldine Conejo-Barboza, Raul Mora-Amador, Keilor Rojas, Dietmar H. Pieper, Max Chavarría

## Abstract

In this work, we characterize the geochemistry and microbial community of Bajo las Peñas, a neutral (pH 6.5-7.4), hot spring (T = 62.0-68.0°C) located near Turrialba Volcano, Costa Rica. The microbiota at its two sources belongs mainly to the family Aquificae, comprising OTUs closely related to the genera *Sulfurihydrogenibium*, *Thermosulfidibacter*, *Thermodesulfovibrio* and *Thermocrinis* which is consistent with the presence of moderate levels of sulfate (243-284 mg/L) along the stream. We determined a dramatic shift in the microbial community just a few meters downstream of the sources of the hot spring (15-20 meters), with a change from sulfur related chemoautotrophic (e.g. *Sulfurihydrogenibium* and an OTU closely related to *Thermodesulfovibrio*) to chemoheterotrophic prokaryotes (e.g. *Meiothermus*, *Nitrososphaera*, *Thermoflexus*, *Thermus*). Thus, in this neutral hot spring, the first level of the trophic chain is associated with photosynthesis as well other anaerobic CO_2_ fixing bacteria. Then, thermotolerant chemoheterotrophic bacteria colonize the environment to degrade organic matter and use fermentative products from the first level of the trophic chain. Our data demonstrate how quickly the microbial community of an ecosystem can change in response to environmental variables and sheds light on the microbial ecology of less common circumneutral pH hot springs.

## Introduction

Characterization of microbial communities inhabiting hot springs is of great interest for understanding the ecology in extreme environments. Due to their high temperatures, macroscopic organisms are limited in these habitats; as a result, hot spring communities are usually made up exclusively by thermotolerant or thermophilic microorganisms (Akanbi et al. 2019). These conditions define hot springs as “ecologic islands”, where even when foreign microorganisms are constantly inoculated into the water, they are not able to survive the high temperatures. Additionally, due to their geological and chemical properties, a great spectrum of niches exists in this kind of environments, which in consequence allows the coexistence of multiple species (Mitri 2019). The great diversity present in the multiple niches (microbial mat, sediments, surface, and underground water) has been proposed to be a consequence of metabolic heterogeneity that evokes during the community evolution (Mitri 2019). Such a variation of taxa across different niches of the same hot spring has been observed when analyzing the microbial mat and the anoxic region of Mushroom Spring in Yellowstone National Park (Klatt et al. 2011; Thiel et al. 2016; Tank et al 2017). Furthermore, in this hot habitat, it has been shown that specific taxa are associated to particular metabolic functions within the community and cross-feeding may be occurring explaining the high diversity in these habitats (Thiel et al. 2017). The understanding of the rules that govern the ecology of hot springs is relevant because these extreme habitats have also been proposed to be a good model for understanding the evolution of life, particularly because hot springs resemble hydrothermal vents where life might have emerged (Damer and Deamer 2020).

Hot springs can be found at various locations around the globe and their physicochemical parameters vary widely but they are normally acidic. Previous studies have demonstrated that their acidity levels are highly correlated to the chemical species that are present in the environment, particularly iron and sulfur species (Xu et al. 1998, 2000; Pierson et al. 1999, 2000; Sharma et al. 2001). A particularity shared between the less commonly described neutral pH hot springs with the acidic ones is the significant proportion of ferrous iron produced from mineral dissolution, which can form precipitates (Konhauser 1997; Pierson et al. 1999; Yee et al. 2003). The iron minerals in neutral hot springs are usually formed by spontaneous iron oxidation, and the oxides produced consequently nucleate, sometimes in association with silica, and precipitate as ferric hydroxides that include minerals like ferrihydrite, lepidocrocite or goethite (Peng et al. 2013; Liu et al. 2014; Cooper et al. 2017). Due to their mild pH and moderate toxic metals concentrations, the main extreme conditioning factor in neutral hot springs is temperature, limiting the proliferation of multiple microorganisms and directly shaping the community (Ward et al. 1998).

Photoautotrophic microorganisms play a key role in hot water bodies (Ward and Castenholz 2000; Klatt et al. 2011). Previous reports have found that members of Cyanobacteria and Chloroflexi are common phototrophic groups found in this type of habitats, particularly in microbial mats (Pierson and Castenholz 1974; Miller and Castenholz 2000; Alcamán et al. 2015; Prieto-Barajas et al. 2018; Ward et al. 2018). On the other hand, some deep sulfurous hot springs have a low abundance of cyanobacteria but a high concentration of sulfur chemoautotrophic bacteria, particularly from the family Desulfobacteriaceae. In these environments, these microorganisms are the main CO_2_-fixers (Nunoura 2014). In addition to the importance of autotrophic microorganisms, multiple taxa classified as heterotroph, such as Deinococcus-Thermus, Nitrospirae, Proteobacteria, Firmicutes and Actinobacteria are also an important part of hot spring communities (Valverde et al. 2012; Wang et al. 2013; Daims 2014; Sahoo et al. 2015; Ghilamicael et al. 2017; Rozanov et al. 2017). Complete metagenome studies have also revealed that these microbial communities can have an extremely diverse and complex metabolism, where nutrients can be recycled between phylotypes and multiple cross-feeding processes may be occurring (Mangrola et al. 2015; Saxena et al. 2017; Thiel et al. 2017; Pedron et al. 2019).

Bajo las Peñas is a natural occurring hot spring located in Cartago province, close to Turrialba and Irazu volcanoes (Fig. 1). This site is a small lotic environment where water from the hot spring source flows slowly through an extensive bacterial mat for about 20 meters and then joins a cold, mountain stream that eventually becomes a tributary of the Toro Amarillo River. It is characterized for having a reddish gelatinous microbial mat covering the site (Figs 1C and 1D), a neutral pH (6.5-7.4) and a temperature of around 65°C (Uribe‐Lorío et al. 2019). Study of circumneutral hot springs like Bajo las Peñas is relevant since they are uncommon in the world (most are acidic) and can potentially become a source of microorganisms and thermostable enzymes of biotechnological interest. After we carried out our sampling in 2016 for this study, a survey of the microbiology of several Costa Rican hot springs reported data of the microbial community inhabiting the microbial mat of Bajo Las Peñas based on samples taken in 2012. (Uribe‐Lorío et al. 2019). This last study reported that the microbial community in the mat is formed almost exclusively by Cyanobacteria. Here, we show a more extensive study of the geochemistry and microbial communities that inhabit the water and the sediments in Bajo las Peñas, including samples taken right at the source in order to better understand the microbial diversity and the possible metabolism associated with this close-to-neutral pH hot spring.

**Fig. 1.**
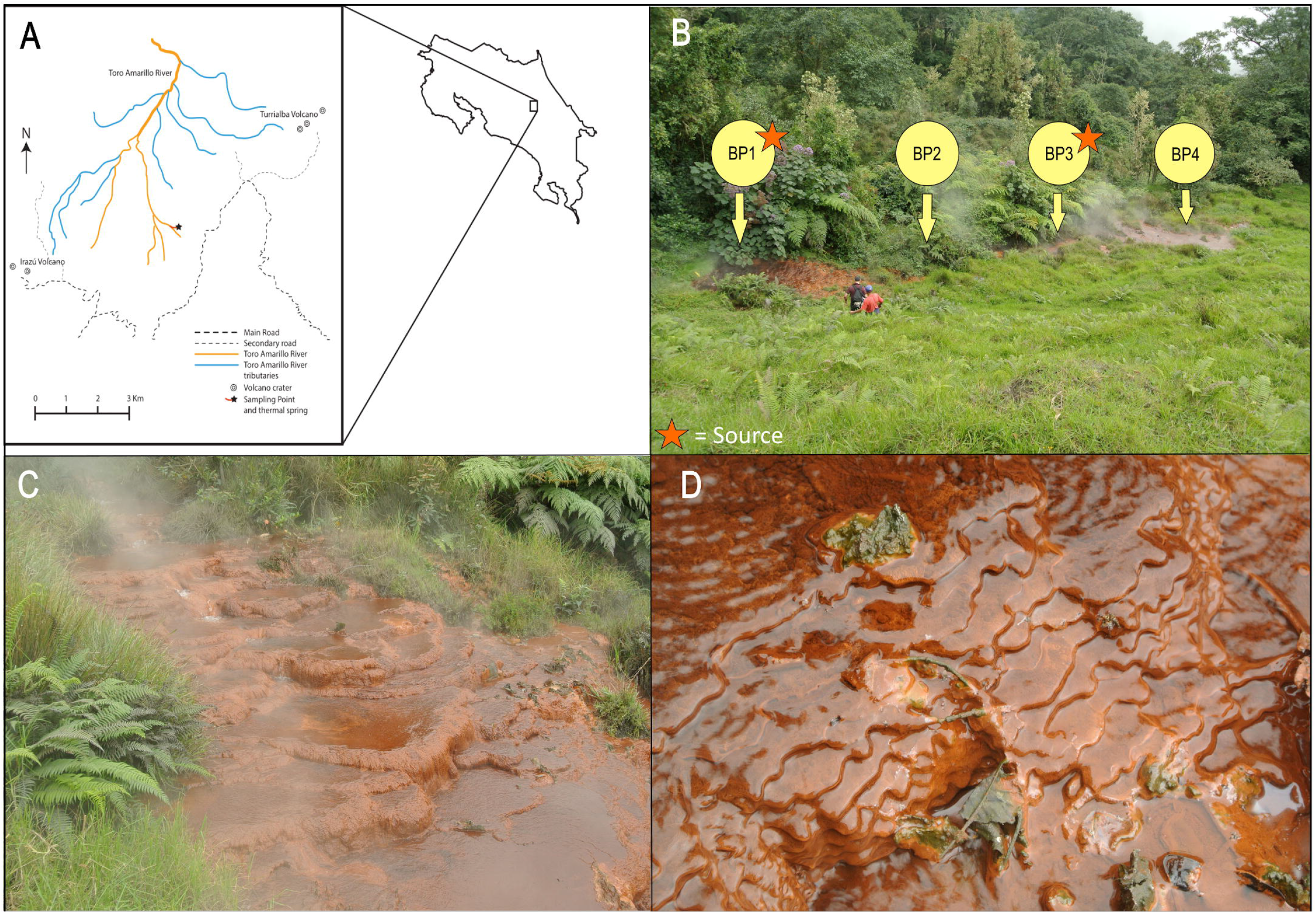
Bajo las Peñas hot spring. A) Bajo las Peñas hot spring is located in Cartago Costa Rica near Turrialba Volcano and Toro Amarillo River. B) It is a small lotic hot spring, with two springs (BP1 and BP3) where water emerges from underground deposits. C) The water flow around Bajo las Peñas is very slow and it flows for about 20 meters. D) The entire hot spring is covered by a dense reddish gelatinous microbial mat where ferryhydrite precipitate constantly.

## Materials and Methods

### Study Site

Bajo las Peñas is located in a private farm northeast of Irazu Volcano (Cartago, Costa Rica; Fig. 1A). At the site, we actually identified and sampled two springs (sources) where hot water emerges which are separated by about 15 m (BP1 and BP3; see Fig. 1). The samples were taken along the whole site, which is approximately 20 meters long and included two more sampling sites away from the sources. Along the effluent, a very thick (ca. 30-50 cm) reddish microbial mat covers the site and slows down the flow of water. Interestingly, despite the high temperature in the surroundings of this thermal spring, grass and bushes were found. The BP1 point (Fig. 1; 9.993492 N, 83.801597 W) corresponds to the origin (first spring) of Bajo las Peñas. The second sampling point (BP2; 9.993558 N 83.801639 W) is located 10 m downstream. Five meters further downstream of BP2 (15 m from the first spring) emerges the second spring, which corresponds to our third sampling point (BP3; 9.993603 N, 83.801677 W). Finally, 5 m downstream from BP3 (20 meters downstream from BP1), our fourth sampling point (BP4; 9.993658 N 83.801716 W) was selected, just a few meters before Bajo las Peñas mixes with a cold stream. Permits for sampling were obtained from the Institutional Commission of Biodiversity of the University of Costa Rica (resolution N° 066) and the owners of the property.

### Sampling and field measurements

On 18 August 2016, samples of water (three samples of 1 Liter each) were collected at four points (BP1 to BP4; see Fig. 1) along this thermal water (i.e. twelve water samples). For this purpose, sterile flasks were attached to a long rod in order to reach down into the water and avoid contamination. At each sampling point it was possible to take one or two sediment samples, for a total of seven sediment samples across the entire spring. Specifically, two sediment samples were taken from sampling site 1 (BP1S1 and BP1S2), two from sites 2 (BP2S1 and BP2S2) and 4 (BP4S1 and BP4S2), whereas a single sediment (BP3S1) was retrieved from point 3. For sediment sampling, masses between 10 and 50 g were placed into sterile 50 mL centrifuge tubes. The sediments were taken at depths not exceeding 15 cm and due to the presence of the microbial mat, the sediment samples contain part of it. In the field, temperature was measured with a partial immersion thermometer. To measure dissolved oxygen (DO), bottles (1 L) were filled with water from each sampling point, hermetically sealed and cooled to room temperature (~22 °C). At that temperature the dissolved oxygen was measured with a dissolved oxygen meter Model 550A (Yellow Springs Instrument Company Inc, Ohio, USA). pH was measured with a pH meter (Scholar 425 pH meter, Corning, Inc., Corning, NY) in the laboratory. Water samples for chemical analysis were collected in clean glass bottles, chilled on ice, and stored at 4 °C until analysis. Water samples for microbial community analysis were collected in clean and sterile bottles, stored at 4 °C and processed within 24 h.

### Chemical Analysis

For chemical analyses, water samples were filtered through polycarbonate membrane filters (0.45 μm; Sartorius 23006-47N) before analysis. Major anionic components (Cl-, F-, NO_3_-, SO_4_^-2^) in the water samples were analyzed by ion-exchange chromatography (IC, MIC-II, Metrohm Co., Switzerland) using an anionic exchange resin (Metrosep A Supp 5 - 100/4.0). Operating conditions were a mobile phase at 33 °C, Na_2_CO_3_ (3.2 mM) / NaHCO_3_ (1.0 mM) and flow rate 0.7 mL/min. The anions were identified and quantified relative to certified commercial standards (Certipur ®Anion multi-element standard I, II, Merck, Germany). On the other hand, elemental components (Al, As, Cd, Ca, Cu, Cr, Fe, Mg, Mn, Ni, Pb, K, Na) were analyzed by inductively-coupled-plasma mass spectrometry (ICP-MS, Agilent 7500 instrument, Agilent Technologies, Tokyo, Japan) using a certified multi-element stock solution (Perkin-Elmer Pure Plus standard, product number 9300233). All determinations were made with three independent samples and the reported results correspond to their average. The uncertainty (±) was calculated using the standard deviation (s) of the three repetitions (n = 3) with a coverage factor K = 2.

### X-Ray Diffraction (XRD) analysis

Powder X-ray diffraction patterns were recorded with a D8 Advance diffractometer (Bruker AXS, Madison, USA, LynxEye detector) with Cu radiation (*λ* = 0.154 nm) from 2θ 15° to 80° and step size 0.018°. Identification was made with ICDD PDF-2 2007 Database [ICDD 2007] and comparison with published ferrihydrite diffraction patterns.

### Scanning Electron Microscope and Electron Dispersive Spectrometer (SEM-EDS)

The chemical elements that constitute the reddish mat and sediments were determined with SEM-EDS. For that purpose, samples were analyzed with a scanning electron microscope (Hitachi S-570) with energy-dispersive X-ray spectra (SEM-EDS).

### Total DNA isolation, construction of 16S rRNA gene libraries and Illumina sequencing

The three water samples from each sampling point (3 x 1L each) were pooled and filtered through a vac-uum system under sterile conditions using a membrane filter (pore size 0.22 μm; Millipore, GV CAT No GVWP04700). To prevent rupture, another filter membrane (pore size 0.45 μm; Phenex, Nylon Part No AF0-0504) was placed below. The upper filter was collected and stored at −80 °C until processing. The DNA was extracted from aseptically cut pieces of the filter with a DNA isolation kit (PowerSoil®, MoBio, Carlsbad, CA, USA) as described by the manufacturer. Cell lysis was accomplished by two steps of bead beating (FastPrep-24, MP Biomedicals, Santa Ana, CA, USA) for 30 s at 5.5 m s−1. To process the sediments, a homogeneous sample of 500 mg was collected and DNA was extracted using the same protocol. For the construction of microbial 16S rRNA amplicon libraries, the V5-V6 hypervariable regions were PCR-amplified with universal primers 807F (5’-GGATTAGATACCCBRGTAGTC-3’) and 1050R (5’-AGYTGDCGACRRCCRTGCA-3’) (Bohorquez et al. 2012). The barcoding of the DNA amplicons and the addition of Illumina adaptors were conducted by PCR from approximately 1-10 ng of total DNA as described previously (Burbach et al. 2016). The PCR-amplified amplicons were verified by 1% agarose gel electrophoresis, purified using the QIAquick Gel Extraction Kit (Qiagen, Hilden, Germany) and quantified with the Quant‐iT PicoGreen dsDNA reagent kit (Invitrogen, Darmstadt, Germany). The individual amplicons were pooled to equimolar ratios (200 ng of each sample) and purified with the QIAquick PCR Purification Kit (Qiagen, Hilden, Germany). Finally, PCR-generated amplicon libraries were subjected to 250 nt paired-end sequencing on a MiSeq platform (Illumina, San Diego, CA, USA).

### Bioinformatic and phylogenetic analysis of 16S rDNA amplicon data

The sequence data were deposited in the sequence-read archive (SRA) of GenBank under the BioProject PRJNA678689. Bioinformatic processing was performed as previously described (Schulz et al. 2018). Raw reads were merged with the Ribosomal Database Project (RDP) assembler (Cole et al. 2014). Sequences were aligned within MOTHUR (gotoh algorithm using the SILVA reference database; (Schloss et al. 2009) and subjected to preclustering (diffs=2) yielding the so-called operational taxonomic units (OTUs) that were filtered for an average abundance of ≥0.001% and a sequence length ≥250 bp before analysis. OTUs were taxonomically classified based on the SILVA v138 database with the RDP taxonomy (Yilmaz et al. 2014) as reported by the SINA classification tool (Pruesse et al. 2012). OTUs were assigned to a taxonomic rank only if their best hit in the SILVA database (Pruesse et al. 2007) had an identity higher than the threshold established by Yarza et al. (2014) for that rank (94.5% for genus, 86.5% for family, 82.0% for order, 78.5% for class and 75.0% for phylum). Moreover, the sequences of some highly abundant OTUs were also manually examined by means of BLASTN (Altschul et al. 1997) against the non-redundant and against the bacterial and archaeal 16S rRNA databases. Sequences that weren’t assigned to at least a phylum were excluded for further analysis. The statistical analyses and their visualization were performed with the R statistical program (R-Core-Team 2019) and Rstudio interface. Package Vegan v2.5-6 (Oksanen et al. 2019) was used to calculate alpha diversity estimators (Observed richness, Shannon index, Simpson index). Data tables with the OTUs abundances were normalized into relative abundances and then converted into a Bray–Curtis similarity matrix. The effects of factors on the bacterial community composition were evaluated with non-parametric permutational analysis of variance (PERMANOVA) (adonis2 function with 999 permutations) and using multivariate analysis such as Principal Coordinate Analysis (PcoA). Significant differences between a priori predefined groups of samples were evaluated using Permutational Multivariate Analysis of Variance (PERMANOVA), allowing for type III (partial) sums of squares, fixed effects sum to zero for mixed terms. Monte Carlo p-values (p_MC_) were generated using unrestricted permutation of raw data (Anderson 2001) IN PRIMER (v.7.0.11, PRIMER-E, Plymouth Marine Laboratory, UK). We used the Kruskal Wallis test to determine significant differences in alpha diversity estimators (Simpson and Shannon-index).

## Results and Discussion

### Chemical Analysis

The major physicochemical properties of Bajo las Peñas are shown in Table 1. This habitat corresponds to a hot spring with moderately high temperature, ranging from 62.0-68.0°C, and pH nearly neutral, ranging from 6.5 to 7.4. Despite the small differences in pH and temperature throughout the site, we observed small but consistent differences between BP1 and BP3 (the two sources) and the stream sites BP2 and BP4 being the sources slightly more acidic and hotter. The oxygen levels (2.01-2.53 mg O_2_/L measured at 22°C) in all the samples indicate that Bajo las Peñas corresponds to a dysoxic environment. (Spietz et al. 2015). Physicochemical analysis of water samples showed the presence of moderate levels of sulfate (243-284 mg/L), magnesium, manganese, sodium, aluminum, calcium, iron and potassium. Our data (samples taken in 2016) agree with the chemical composition reported by Uribe-Lorío et al (2019) of samples taken in 2012, suggesting that the inorganic chemistry of Bajo Las Peñas has remained relatively stable. Examination of the dried sediment samples revealed considerable heterogeneity and organic matter content. Even a few *ca*. 0.2 mm sized magnetite and sand grains were present and were removed prior to the X-Ray measurements. This heterogeneity is expected as the sediments do not consist of a pure mineral. EDS data showed sediments to be iron-rich though extremely variable (50-70% Fe) and we assign them to ferric hydroxide precipitates. The samples were very amorphous, required several hours of Powder X-Ray data collection and only one did diffract strongly enough to identify a mineral phase. The XRD pattern (Fig. S1) of the reddish sediment sample from site BP1 was assigned to 6-line ferrihydrite rather than the less crystalline 2-line ferrihydrite. The spacings for the diffraction peaks at 2.53, 2.22, 1.96, 1.72 1.50 and 1.46 Å compare very well to the literature (Fig. S1; Drits et al. 1993). Only an unassigned peak at 2.44 Å remained. Interestingly, the conversion from 2-line ferrihydrite into 6-line ferrihydrite has been reported to occur under biotic conditions (Kukkadapu et al. 2003). Ferrihydrite is normally very amorphous and the presence of organic matter made collecting Powder X-Ray data difficult. We could not obtain a diffraction pattern from other sediment samples, and we include as an example the essentially absent diffraction of the sample from the BP2 site as reference (Fig. S1). Additionally, previous reports of similar environments indicate that the mixture of the sediments and the microbial mat with the water flow produces the characteristic red color of these habitats (Casanova et al. 1999; Bishop and Murad 2002; Naren et al. 2012), as is the case in Bajo las Peñas (see Fig. 2).

**Table 1.** Physical properties and chemical composition of Bajo las Peñas hot spring.

| Property/element/ion | BP-1 | BP-2 | BP-3 | BP-4 |
| --- | --- | --- | --- | --- |
| Coordinates | 9.993492 N<br>83.801597 W | 9.993558 N<br>83.801639 W | 9.993603 N<br>83.801677 W | 9.993658 N<br>83.801716 W |
| Distance from the first origin (m) | 0 | 10 | 15 | 20 |
| Temperature (°C) / $\pm 0.1$ | 68.0 | 65.6 | 67.0 | 62.0 |
| pH / $\pm 0.1$ | 6.5 | 7.4 | 6.6 | 7.3 |
| Dissolved oxygen (measured to 22°C) | 2.01 | 2.05 | 2.23 | 2.53 |
| Aluminium ($\mu\text{g L}^{-1}$) $\pm 3$ | 15 | 25 | 13 | 14 |
| Arsenic ($\mu\text{g L}^{-1}$) | <2 | <2 | <2 | <3 |
| Cadmium ($\mu\text{g L}^{-1}$) | < 0.11 | < 0.17 | < 0.12 | < 0.11 |
| Calcium ($\text{mg L}^{-1}$) $\pm 2$ | 15 | 11 | 20 | 11 |
| Chloride ($\text{mg L}^{-1}$) $\pm 1$ | < 0.20 | 1 | 1 | 1 |
| Cooper ($\text{mg L}^{-1}$) | < 0.10 | < 0.10 | < 0.10 | < 0.10 |
| Chromium ($\mu\text{g L}^{-1}$) | < 1.2 | < 1.2 | < 1.2 | < 1.2 |
| Fluoride ($\text{mg L}^{-1}$) | <0.09 | <0.09 | <0.09 | <0.09 |
| Iron ($\text{mg L}^{-1}$) / $\pm 0.06$ | 0.60 | 0.16 | 0.16 | <0.08 |
| Magnesium ($\text{mg L}^{-1}$) $\pm 0.7$ | 11.6 | 12.2 | 12.6 | 13.0 |
| Manganese ($\text{mg L}^{-1}$) $\pm 0.06$ | 0.36 | 0.36 | 0.41 | 0.40 |
| Nickel ($\mu\text{g L}^{-1}$) | < 1 | < 1 | < 1 | < 1 |
| Nitrate ($\text{mg L}^{-1}$) | < 0.10 | < 0.10 | < 0.10 | < 0.10 |
| Lead ($\mu\text{g L}^{-1}$) | < 1.2 | < 1.2 | < 1.2 | < 1.2 |
| Potassium ($\text{mg L}^{-1}$) $\pm 0.3$ | 9.8 | 10.0 | 10.6 | 10.6 |
| Sodium ($\text{mg L}^{-1}$) $\pm 0.8$ | 38.5 | 39.3 | 38.1 | 38.5 |
| Sulfate ($\text{mg L}^{-1}$) $\pm 20$ | 264 | 281 | 284 | 243 |
| Zinc ($\mu\text{g L}^{-1}$) | < 25 | < 25 | < 25 | < 25 |

**Fig. 2.**
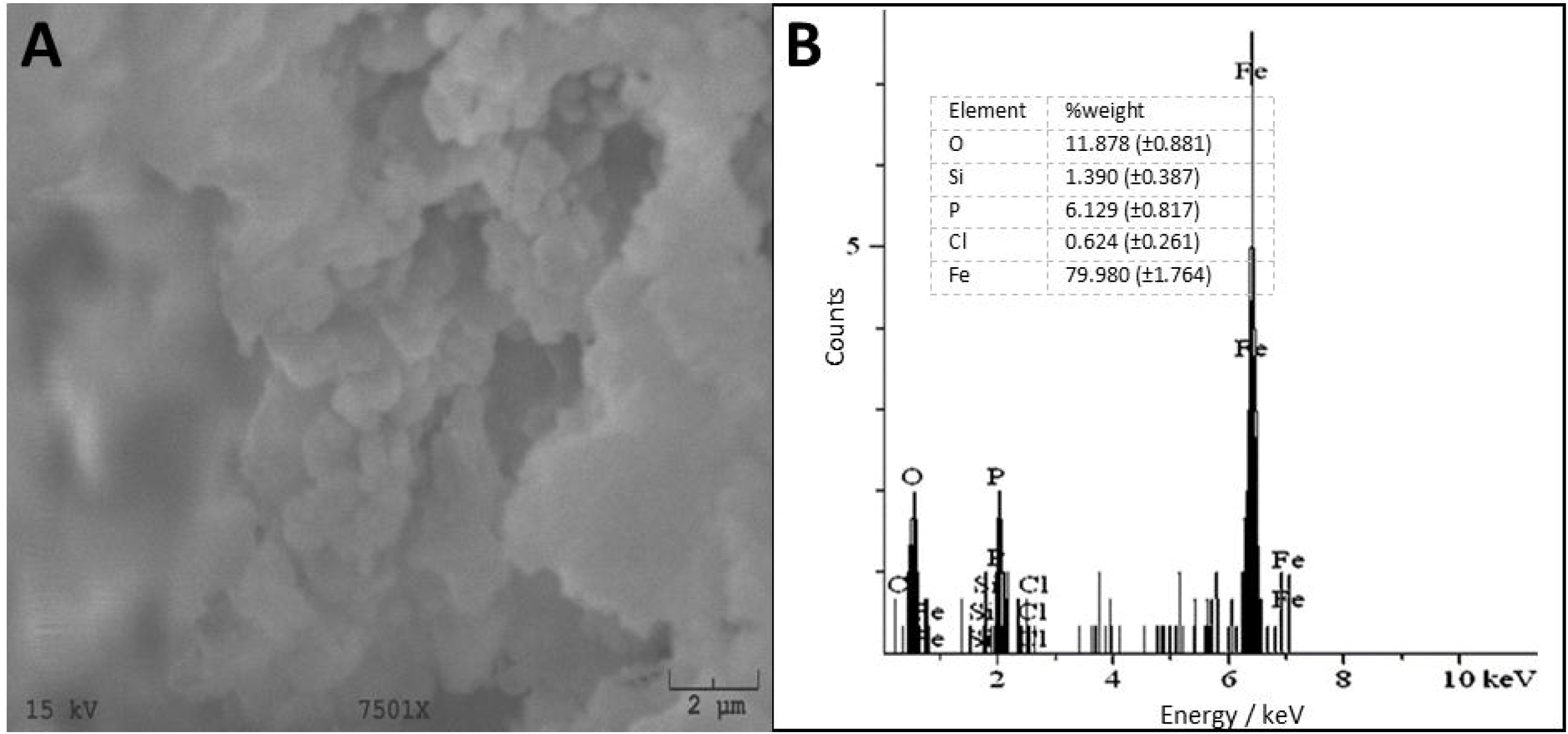
SEM-EDS analysis of Bajo las Peñas sediments. A) SEM micrographs to 2 μm (15.0 kV 7.6 mm × 60) of the reddish gelatinous sediments at the bottom of Bajo las Peñas. Micrographs indicate that the sediment consists of lumps of mainly amorphous material of nanometer size. B) EDS analysis of the sediments showed that it is composed of mainly iron and oxygen. Other species such as silicon, phosphate and chlorine were found in the sediment in minor proportions.

In spite that Bajo las Peñas is located close to other acidic rivers (e.g. Rio Sucio and Toro Amarillo rivers) likely associated with the area’s sulfate and chloride-rich hydrothermal water, Bajo las Peñas physicochemical properties are particularly different: it is endowed with moderate sulfate concentrations in the water column, low chloride, neutral pH and precipitation of ferrihydrite rather than ferric-oxide sulfate minerals. All these observations suggest that the sulfate source in Bajo las Peñas is different from Acid Rock Drainage (ARD) and Volcanic influenced Acid Rock Drainage (VARD) environments. We have described VARD ecosystems associated with nearby Irazu Volcano’s hydrothermal system (Arce-Rodríguez et al. 2020) in which the sulfate and the acidity come from the volcano hydrothermal system rather than from biological oxidation of pyrite as in ARD systems (Amils et al. 2002; Arce-Rodríguez et al. 2017, 2019). However, they do not seem to influence Bajo las Peñas and instead, the likely sources of sulfate are anhydrite/gypsum dissolution perhaps aided by some acid rain as the area is downwind from active Turrialba Volcano (Porowski et al. 2019). The hydrothermal system at Bajo las Peñas contributes therefore mostly heat and ferrous iron to the aquifer rather than acid-forming magmatic gases.

### Analysis of microbial communities

We determined the presence of 916 OTUs in all samples, which were taxonomically assigned to 35 phyla of Bacteria and Archaea (Supplementary Table 1). The most-abundant phyla were Actinobacteria (0.8-40.0%), Proteobacteria (4.9-39.7%), Chloroflexi (0.8-29.1%), Nitrospirae (2.6-28.6%), Firmicutes (0.4-25.9%), Thaumarchaeota (0.1-25.8%), Crenarchaeota (0.1-24.0%), Deinococcus-Thermus (0.4-19.6%), Aquificae (0-16.0%), Ignavibacteriae (0-14.0%), Acetothermia (0-5.6%), Dictyoglomi (0-5.1%), Acidobacteria (0.8-4.7%), Cyanobacteria (0-4.4%), and Planctomycetes (0-4.0%). Nonetheless, we observed large variations across samples (Fig. 3).

**Fig. 3.**
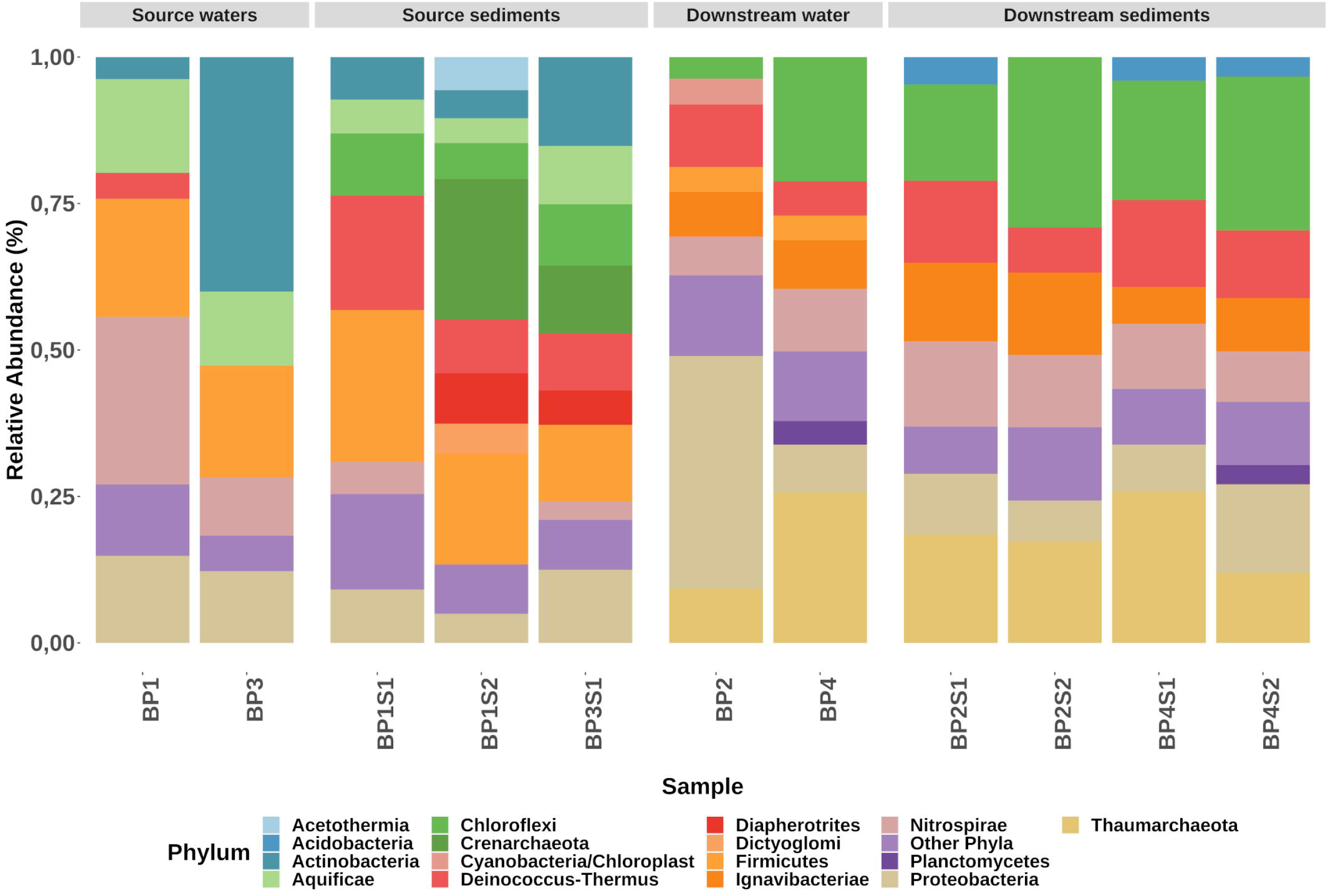
Taxonomic composition of Bajo las Peñas microbial communities. Relative abundance of bacterial and archaeal organisms to phylum. The OTUs were taxonomically classified using RDP database, as described in Materials and Methods. Waters of Bajo las Peñas are termed BP1 to BP4 and sediments are termed BP1S1 to BP4S2.

Comparisons between the global bacterial assemblages in the different sites showed the presence of three well-defined clusters: (i) water springs (BP1, BP3), (ii) sediments in the springs (BP1S1, BPS2, BP1S3), (iii) waters and sediments downstream from the springs (BP2, BP2S1, BP2S2, BP4, BP4S1, BP4S2) (Fig. 4) (PERMANOVA p = 0.001). All groups of samples were significantly different from one another as indicated by pairwise tests (source water vs downstream samples p_MC_ = 0.05, source sediments vs downstream samples, p_MC_ = 0.01 and source water vs source sediment, p_MC_ = 0.04). This clustering pattern indicates a fast community turnover right after communities emerge from underground to the surface, possibly in response to new environmental conditions such as increase in oxygen and carbon input.

**Fig. 4.**
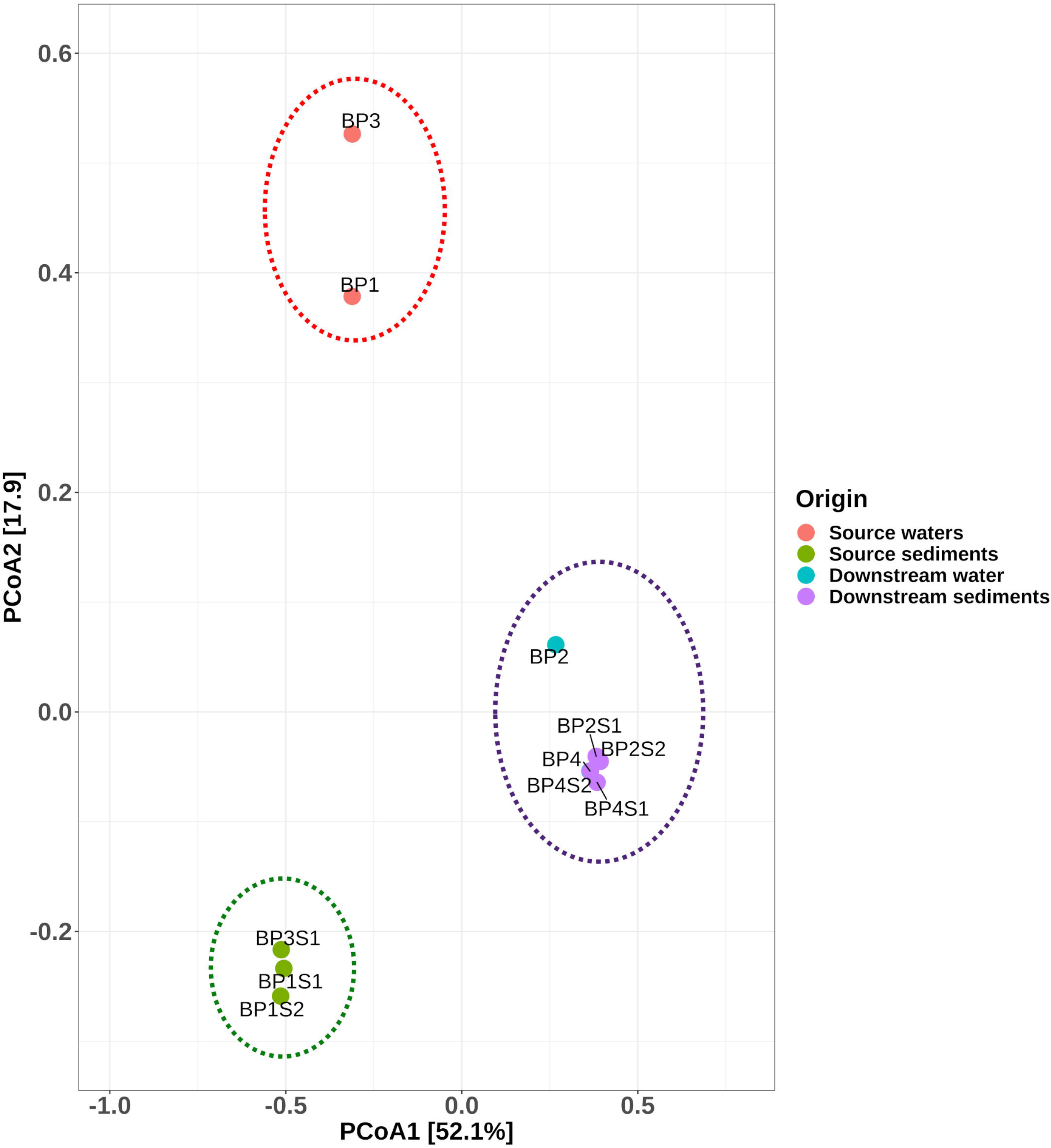
Principal Coordinate Analysis (PCoA) of the prokaryotic communities in Bajo las Peñas. A clustering of communities according to the sampling point is shown. The PCoA and the PERMANOVA analyses were performed with the package Vegan.

Alpha diversity estimations (Fig. S2) showed that all samples present a Shannon index > 2.8. Kruskal-Wallis test showed no significant differences in the diversity indices across the clusters described. Previous analyses of the microbial mats in Bajo las Peñas (Uribe‐Lorío et al. 2019) showed a high abundance of photoautotrophic prokaryotes, where almost the entire the community was dominated by few OTUs. However, our analyses, including samples in the water column and sediments showed that this environment holds a very variable microbial community (See Figs 3 and 4). The high dynamism that exists in this extreme environment, where the community fluctuates according to minor changes in the water conditions and external factors, such as the climate (Ferris and Ward 1997; Fraterrigo et al. 2006; Li et al. 2014; Narayan et al. 2018) is also a factor to consider when comparing results. Interestingly, the absence of iron-oxidizing bacteria (see below) can be related to the neutral pH which favors its abiotic oxidation, thus reducing the availability of electron donors for this type of bacteria (Earhart 2009; Johnson et al. 2012). This is one of the clearest differences in Bajo las Peñas compared to other hot springs that we have characterized in Costa Rica (Arce-Rodríguez et al. 2019, 2020), whose physicochemical characteristics are typical of ARD or VARD environments (with acidic pH), where the presence of iron oxidizers is very common.

The most abundant microorganisms in the origins of Bajo las Peñas (water samples BP1 and BP3) were phylotypes from the genus **Sulfurihydrogenibium**, particularly OTU_439 (10.3-14.8%) (Fig. 5) and some putative members of the family Nitrospiraceae (OTU_67, 10.0-26.8%), which closest relatives were bacteria of the genus **Thermodesulfovibrio** (92.2% identity). The genus **Sulfurihydrogenibium** has to our knowledge five reported species: *S. azorense* (Aguiar et al. 2004), *S. kristjanssonii* (Flores et al. 2008), *S. rodmanii* (O’Neill et al. 2008), *S. subterraneum* (Takai et al. 2003), and *S. yellowstonense* (Nakagawa et al. 2005). These microorganisms are reported as aerobic, thermophilic, and sulfur-oxidizing bacteria. These bacteria isolated from hot springs grow at temperatures between 40-80°C and can use S°, thiosulfate and sulfite as electron donors. Some species from this genus have reported the ability to use CO_2_ as carbon source. Moreover, these microorganisms are known to be important organic carbon producers in the environments where they were isolated (Nunoura 2014). Thus, we suggest that members of the genus **Sulfurihydrogenibium** could act as CO_2_-fixers in Bajo Las Peñas.

**Fig. 5.**
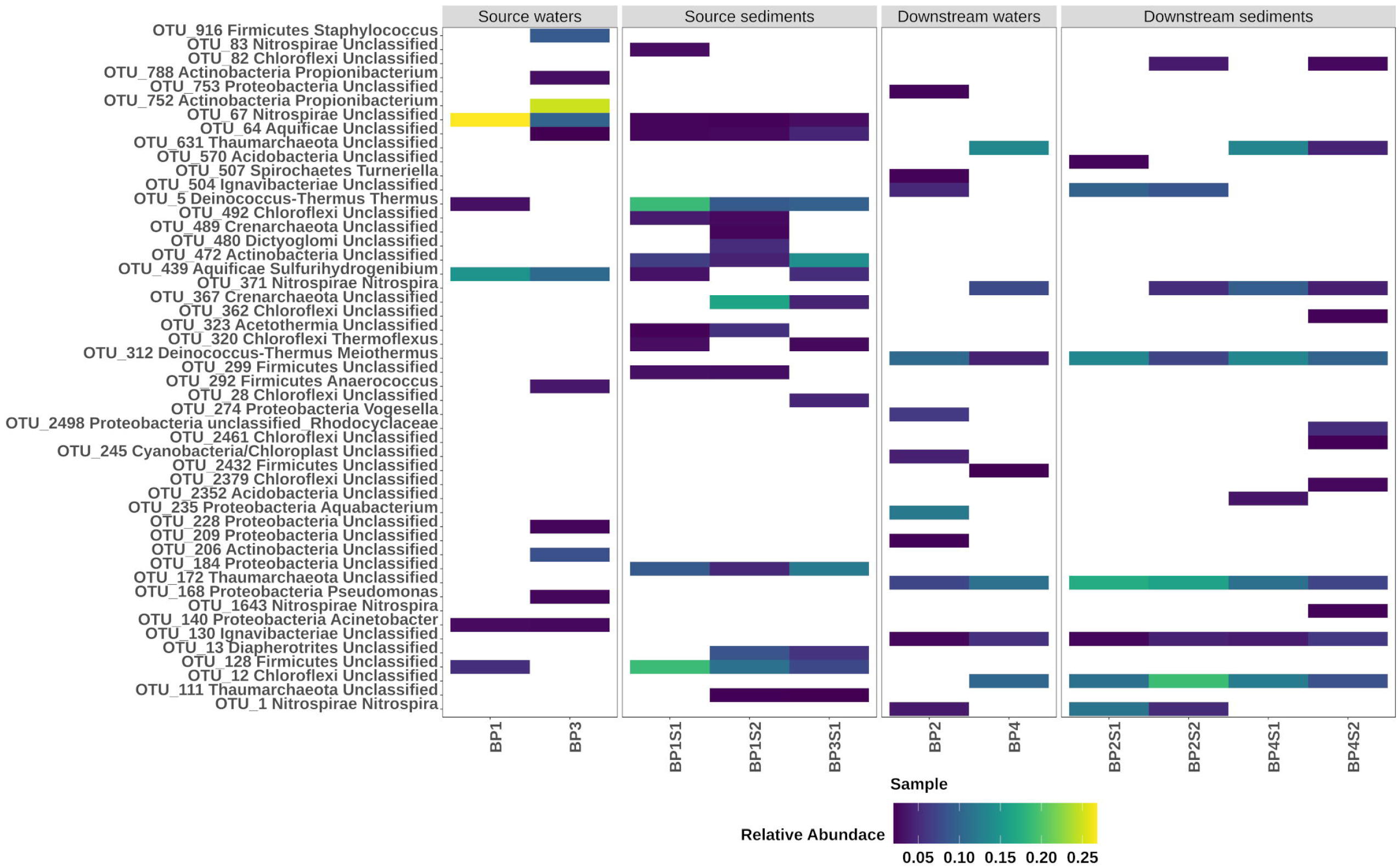
Heat map of the most abundant OTU in each sample. The heat map depicts the relative percentage of 16S rRNA gene sequences assigned to each OTU (y axis) across the 11 samples analyzed (x axis) with an abundance higher than 2%.

On the other hand, the relatively low sequence identity of OTU_67 with members of the genus *Thermodesulfovibrio*, does not allow us to reliably affirm that they belong to this genus. However, considering that (i) they belong to the family Nitrospiraceae, (ii) the moderate levels of available sulfate, and (iii) the temperature in Bajo las Peñas, presumably members of OTU_67 could participate in sulfate reduction in a similar way as species of *Thermodesulfovibrio* genus do. Bacteria from the *Thermodesulfovibrio* genus have typically been isolated from hot springs and are generally known as anaerobic sulfate reducers that grow at temperatures between 40-70°C. *T. yellowstonii* has the ability to use sulfate, thiosulfate, sulfite and nitrate as terminal electron acceptors and oxidizes organic compounds like pyruvate and lactate to acetate. Autothrophic growth using hydrogen have also been reported in members of this genus (Daims 2014).

In the sediments taken from Bajo las Peñas sources it was possible to find the same OTU from the genus *Sulfurihydrogenibium* (OTU_439, 1.6-5.2%) and the OTU_67 (2.2-2.8%) found mostly in the waters. However, many other OTUs closely affiliated to *Thermoflexus* (OTU_320, 1.9-2.9%) and Thermus (OTU_5, 8.9-18.9%), to the families Dictyoglomaceae, Thermomicrobiaceae and Aquificaceae respectively, (OTU_480, 1.9-5.1%; OTU_492, 2.7-3.8% and OTU_64, 2.5-4.6 %) or to the Clostridia (OTU_128, 7.5-18.9%) were also detected with a high abundance (Figure 5).

Bacteria from the genera *Thermoflexus* and *Thermus* have been reported to be chemoheterotrophic microorganisms, that may consume byproducts of other microorganism or substrates deposited in the sediments of Bajo las Peñas. Bacteria of the phylum Chloroflexi were also detected in some of these sediment samples (6.1-10.6%) (Fig. 5), which is in accordance with a report by Uribe-Lorío et al. (2019) on the microbial mat. However, our data suggest that in Bajo las Peñas the first level of the trophic chain is not only associated with photosynthesis by Cyanobacteria and Chloroflexi but also by CO_2_ fixation linked to sulfur metabolism by autotrophic bacteria which derives probably from the subdeposits of water in Bajo las Peñas.

A few meters downstream of the origin (BP2 and BP4) the organics produced in the first trophic level (described above) as well as the organic matter input may generate the conditions necessary for thermotolerant chemoheterotrophic bacteria to colonize the place. This would a drastic shift in the microbial community structure with respect to the samples obtained at the origin, although spatially the samples are separated by only 5-20 meters. In the water samples BP2 and BP4, bacteria of the family Nitrososphaera (8.2-25.8%), Ignavibacteriaceae (7.3-7.9%) and of the genus *Nitrospira* (6.2-9.7%) and *Meiothermus* (5.6-10.6%) were dominating. Most of these phylotypes were also dominating in the sediment samples from sampling points 2 and 4 (see Fig. 5). We propose that many of these microorganisms found downstream the origins (sampling points 2 and 4), use fermentation products generated by other microorganisms found at the origin (sampling points 1 and 3). For instance, fermentation products like acetate, have been reported to be produced by members of the genera *Thermodesulfovibrio* and *Ignavibacterium* (Lino et al. 2009) which were closely related to the microorganism identified in our samples. These byproducts could be used as a carbon source by some acetate-oxidizing bacteria. These observations suggest a continuous cross-feeding between the different niches of this hot spring.

Besides the possible carbon cross-feeding, a well-characterized relationship between ammonium-oxidizing and nitrite-oxidizing microorganisms may be occurring. Members of the family Nitrososphaera (2.1-25.7%), may oxidize ammonium to nitrite, which is subsequently utilized by nitrite-oxidizing Nitrospira (6.2-14.0%), for growth (Hovanec et al. 1998; Stieglmeier et al. 2014; Daims et al. 2015; Lehtovirta-Morley 2018). Such a nitrification process has been reported in other hot springs and is known to be an essential process for the maintenance of the community (Reigstad et al. 2008). Thus, our data are aligned with the existence of multiple cross-feeding processes, which are probably the product of microbial adaptation to the low abundance of organic carbon producers in the environment.

As mentioned above, Bajo las Peñas hot spring is characterized for having a red-gelatinous microbial tappet (Fig. 1C) which, we propose, is made of a complex bacterial community in association with ferrihydrite minerals. Uribe-Lorío et al. (2019) reported that the microbial mat in Bajo las Peñas is composed mainly by Cyanobacteria (*Limnothrix, Leptolyngbya, Phormidium* and *Fisherella*) and to a lower extent by *Meiothermus*. We were also able to detect *Meiothermus* which is considered a key microorganism in hot springs due to its ability to form a strong biofilm by specialized structures (Spanevello and Patel 2004; Masurat et al. 2005; Wu et al. 2009; Mori et al. 2012).

## Conclusions

This study sheds light on the ecology of microbial communities of a rare type of hot springs with neutral conditions located in Bajo las Peñas, Costa Rica. In this hot spring, abiotic precipitation of ferric oxyhydroxides in an extensive bacterial mat contribute to the red-colored appearance of the habitat. The analyses of 16S rRNA sequences revealed that Bajo las Peñas harbors a very complex environment where the microbial community changes along the water flow from a chemoautotrophic to a chemolitotrophic metabolism. CO_2_ fixation occurs through photosynthetic microorganisms (Cyanobacteria and Chloreflexi) but also bacteria that fix CO_2_ linked to sulfur metabolism (i.e. *Sulfurihydrogenibium*). In this first autotrophic level, organic molecules are produced that then can be used by heterotrophic microorganisms such as those closely related to *Thermodesulfovibrio* and *Ignavibacterium*. We propose that the microbial community in Bajo Las Peñas may have a complex metabolism due to niche specialization where cross-feeding processes between phylotypes plays an essential role. Our data demonstrate the quick change in the microbial community of an ecosystem in response to environmental variables such as organic matter and oxygen availability. In Bajo las Peñas it was possible to observe how in just ~5 meters (e.g. BP1 vs BP2) the global microbial community and the dominant species change from autotrophic to heterotrophic microorganisms. This work also helps to understand extremophile communities present in neutral hot springs, which are considerably less explored compared to acidic hot springs, and shows that they are an interesting environment for the bioprospecting of microorganisms and extremozymes with promising biotechnological applications.

## Acknowledgements

We thank Carlos Rodriguez of Centro de Investigación en Contaminación Ambiental (CICA) for the help in the chemical analysis. We also are grateful to Solange Voysest and Jose Jimenez for help in the design of some figures.

## Author contributions

AAR, FPS, EL, MC conceived and designed the experiments; AAR, RA, RMA, DRG, PF, GCB performed the experiments; AAR, FPS, DRG, KR, MC analyzed the data; PF, MC, DHP contributed reagents or materials or analytical tools; DRG, EL, KR, DHP, MC wrote the paper. All authors reviewed and approved the final version of the manuscript.

## Compliance with ethical standards

### Ethical approval

This study does not describe any experimental work related to human.

## References

Aguiar P, Beveridge TJ, Reysenbach AL (2004) Sulfurihydrogenibium azorense, sp. nov., a thermophilic hydrogen-oxidizing microaerophile from terrestrial hot springs in the Azores. Int J Syst Evol 54:33–39. https://doi.org/10.1099/ijs.0.02790-0

Akanbi TO, Agyei D, Saari N (2019) Food enzymes from extreme environments: sources and bioprocessing. In: Kuddus M (ed) Enzymes in Food Biotechnology. Elsevier, pp 795–816

Altschul SF, Madden TL, Schäffer AA et al (1997) Gapped BLAST and PSI-BLAST: a new generation of protein database search programs. Nucleic Acids Res 25:3389–3402. https://doi.org/10.1093/nar/25.17.3389

Alcamán ME, Fernandez C, Delgado A, Bergman B, Díez B (2015) The cyanobacterium Mastigocladus fulfills the nitrogen demand of a terrestrial hot spring microbial mat. The ISME Journal 9:2290–2303. https://doi.org/10.1038/ismej.2015.63

Amils R, González-Toril E, Fernández-Remolar D, Gómez F, Rodríguez N, Durán C (2002) Interaction of the sulfur and iron cycles in the Tinto River ecosystem. Rev Environ Sci Biotechnol 1:299–309. https://doi.org/10.1023/a:1023232002312

Anderson MJ (2001) A new method for non-parametric multivariate analysis of variance. Austral Ecol 26:32–46. https://doi.org/10.1111/j.1442-9993.2001.01070.pp.x

Arce-Rodríguez A, Puente-Sánchez F, Avendaño R et al (2017) Pristine but metal-rich Río Sucio (Dirty River) is dominated by Gallionella and other iron-sulfur oxidizing microbes. Extremophiles 21:235–243. https://doi.org/10.1007/s00792-016-0898-7

Arce-Rodríguez A, Puente-Sánchez F, Avendaño R et al (2019) Thermoplasmatales and sulfur-oxidizing bacteria dominate the microbial community at the surface water of a CO2-rich hydrothermal spring located in Tenorio Volcano National Park, Costa Rica. Extremophiles 23:177–187. https://doi.org/10.1007/s00792-018-01072-6

Arce-Rodríguez A, Puente-Sánchez F, Avendaño R et al (2020) Microbial community structure along a horizontal oxygen gradient in a Costa Rican volcanic influenced acid rock drainage System. Microb Ecol 80:793–808. https://doi.org/10.1007/s00248-020-01530-9

Bishop JL, Murad E (2002) Spectroscopic and geochemical analyses of ferrihydrite from springs in Iceland and applications to Mars. Geological Society 202:357–370. https://doi.org/10.1144/gsl.sp.2002.202.01.18

Bohorquez LC, Delgado-Serrano L, López G, et al (2012) In-depth characterization via complementing culture-independent approaches of the microbial community in an acidic hot spring of the Colombian Andes. Microb Ecol 63:103–11. https://doi.org/10.1007/s00248-011-9943-3

Burbach K, Seifert J, Pieper DH, Camarinha-Silva A (2016) Evaluation of DNA extraction kits and phylo-genetic diversity of the porcine gastrointestinal tract based on Illumina sequencing of two hypervariable regions. Microbiologyopen 5:70–82. https://doi.org/10.1002/mbo3.312

Casanova J, Bodénan F, Négrel P, Azaroual M (1999) Microbial control on the precipitation of modern ferrihydrite and carbonate deposits from the Cézallier hydrothermal springs (Massif Central, France). Sedimentary Geology 126:125–145. https://doi.org/10.1016/s0037-0738(99)00036-6

Cole JR, Wang Q, Fish JA et al (2014) Ribosomal database project: data and tools for high throughput rRNA analysis. Nucleic Acids Res 42:633–642. https://doi.org/10.1093/nar/gkt1244

Cooper RE, Eusterhues K, Wegner CE, Totsche KU, Küsel K (2017) Ferrihydrite-associated organic matter (OM) stimulates reduction by Shewanella oneidensis MR-1 and a complex microbial consortia. Biogeosciences 14:5171–5188. https://doi.org/10.5194/bg-14-5171-2017

Core Team R (2017) R: A language and environment for statistical computing. R foundation for statistical computing, Vienna, Austria. http://www.R-project.org/

Daims H (2014) The Family Nitrospiraceae. In: Rosenberg E, DeLong EF, Lory S, Stackebrandt E, Thompson F (eds) The Prokaryotes. Springer, Berlin, pp 733–749

Daims H, Lebedeva EV, Pjevac P et al (2015) Complete nitrification by Nitrospira bacteria. Nature 528:504–509. https://doi.org/10.1038/nature16461

Damer B, Deamer D (2020) The hot spring hypothesis for an origin of life. Astrobiology 20:429–452. https://doi.org/10.1089/ast.2019.2045

Drits VA, Sakharov BA, Salyn AL, Manceau A (1993) Structural model for ferrihydrite. Clay Minerals 28:185–207. https://doi.org/10.1180/claymin.1993.028.2.02

Earhart C F (2009) Iron metabolism. In: Schaechter M (ed) Encyclopedia of microbiology, 3rd edn. Elsevier, pp 210–218

Ferris MJ, Ward DM (1997) Seasonal distributions of dominant 16S rRNA-defined populations in a hot spring microbial mat examined by denaturing gradient gel electrophoresis. Appl Environ Microbiol 63:1375–1381. https://doi.org/10.1128/aem.63.4.1375-1381.1997

Flores GE, Liu Y, Ferrera I, Beveridge TJ, Reysenbach AL (2008) Sulfurihydrogenibium kristjanssonii sp. nov., a hydrogen-and sulfur-oxidizing thermophile isolated from a terrestrial Icelandic hot spring. Int J Syst Evol 58:1153–1158. https://doi.org/10.1099/ijs.0.65570-0

Fraterrigo JM, Balser TC, Turner MG (2006) Microbial community variation and its relationship with nitrogen mineralization in historically altered forests. Ecology 87:570–579. https://doi.org/10.1890/05-0638

Ghilamicael AM, Budambula NLM, Anami SE, Mehari T, Boga HI (2017) Evaluation of prokaryotic diversity of five hot springs in Eritrea. BMC Microbiology. https://doi.org/10.1186/s12866-017-1113-4

Hovanec TA, Taylor LT, Blakis A, Delong EF (1998) Nitrospira-like bacteria associated with nitrite oxidation in freshwater aquaria. Appl Environ Microbiol 64:258–264. https://doi.org/10.1128/aem.64.1.258-264.1998

Johnson DB, Kanao T, Hedrich S (2012) Redox transformations of iron at extremely low pH: fundamental and applied aspects. Front Microbiol. https://doi.org/10.3389/fmicb.2012.00096

Klatt CG, Wood JM, Rusch DB et al (2011) Community ecology of hot spring cyanobacterial mats: predominant populations and their functional potential. The ISME Journal 5:1262–1278. https://doi.org/10.1038/ismej.2011.73

Konhauser KO (1997) Bacterial iron biomineralisation in nature. FEMS Microbiol Rev 20:315–326. https://doi.org/10.1111/j.1574-6976.1997.tb00317.x

Kukkadapu RK, Zachara JM, Fredrickson JK, Smith SC, Dohnalkova AC, Russell CK (2003) Transformation of 2-line ferrihydrite to 6-line ferrihydrite under oxic and anoxic conditions. American Mineralogist 88:1903–1914. https://doi.org/10.2138/am-2003-11-1233

Lehtovirta-Morley LE (2018) Ammonia oxidation: ecology, physiology, biochemistry and why they must all come together. FEMS Microbiol Lett 365(9). https://doi.org/10.1093/femsle/fny058

Li SJ, Hua ZS, Huang LN et al (2014) Microbial communities evolve faster in extreme environments. Scientific Reports 4(6205). https://doi.org/10.1038/srep06205

Lino T, Mori K, Uchino Y, Nakagawa T, Harayama S, Suzuki KI (2009) Ignavibacterium album gen. nov., sp. nov., a moderately thermophilic anaerobic bacterium isolated from microbial mats at a terrestrial hot spring and proposal of Ignavibacteria classis nov., for a novel lineage at the periphery of green sulfur bacteria. Int J Syst Evol 60:1376–1382. https://doi.org/10.1099/ijs.0.012484-0

Liu A, Liu J, Pan B, Zhang W (2014) Formation of lepidocrocite (γ-FeOOH) from oxidation of nanoscale zero-valent iron (nZVI) in oxygenated water. RSC Adv 4:57377–57382. https://doi.org/10.1039/c4ra08988j

Mangrola A, Dudhagara P, Koringa P et al (2015) Deciphering the microbiota of Tuwa hot spring, India using shotgun metagenomic sequencing approach. Genomics Data 4:153–155. https://doi.org/10.1016/j.gdata.2015.04.014

Masurat P, Fru EC, Pedersen K (2005) Identification of Meiothermus as the dominant genus in a storage system for spent nuclear fuel. J Appl Microbiol 98:727–740. https://doi.org/10.1111/j.1365-2672.2004.02519.x

Miller SR, Castenholz RW (2000) Evolution of thermotolerance in hot spring Cyanobacteria of the genus Synechococcus. Appl Environ Microbiol 66:4222–4229. https://doi.org/10.1128/aem.66.10.4222-4229.2000

Mitri S (2019) The evolutionary ecology of microbes. In: Schmidt TM (ed) Reference module in life sciences, 4th edn. Elsevier, pp 416–422

Mori K, Lino T, Ishibashi J, Kimura H, Hamada M, Suzuki K (2012) Meiothermus hypogaeus sp. nov., a moderately thermophilic bacterium isolated from a hot spring. Int J Syst Evol 62:112–117. https://doi.org/10.1099/ijs.0.028654-0

Nakagawa S, Shtaih Z, Banta A, Beveridge TJ, Sako Y, Reysenbach AL (2005) Sulfurihydrogenibium yellowstonense sp. nov., an extremely thermophilic, facultatively heterotrophic, sulfur-oxidizing bacterium from Yellowstone National Park, and emended descriptions of the genus Sulfurihydrogenibium, Sulfurihydrogenibium subterraneum and Sulfurihydrogenibium azorense. Int J Syst Evol 55:2263–2268. https://doi.org/10.1099/ijs.0.63708-0

Narayan A, Patel V, Singh P et al (2018) Response of microbial community structure to seasonal fluctuation on soils of Rann of Kachchh, Gujarat, India: Representing microbial dynamics and functional potential. Ecological Genetics and Genomics 6:22–32. https://doi.org/10.1016/j.egg.2017.11.001

Naren G, Miyazaki A, Matsuo M et al (2012) A study on the interaction between ferric ion and silicic acid in hydrosphere: Si-containing ferruginous deposits formed in neutral hot spring waters. Chinese Journal of Geochemistry 32:27–34. https://doi.org/10.1007/s11631-013-0603-9

Nunoura T (2014) The family Desulfurobacteriaceae. In: Rosenberg E, DeLong EF, Lory S, Stackebrandt E, Thompson F (eds) The Prokaryotes. Springer, Berlin, Heidelberg, pp 617–625

Oksanen J, Blanchet FG, Friendly M et al (2017) Vegan: community ecology package. R package version 2.4–3. https://CRAN.Rproject.org/package=vegan

O’Neill AH, Liu Y, Ferrera I, Beveridge TJ, Reysenbach AL (2008) Sulfurihydrogenibium rodmanii sp. nov., a sulfur-oxidizing chemolithoautotroph from the Uzon Caldera, Kamchatka Peninsula, Russia, and emended description of the genus Sulfurihydrogenibium. Int J Syst Evol 58:1147–1152. https://doi.org/10.1099/ijs.0.65431-0

Pedron R, Esposito A, Bianconi I et al (2019) Genomic and metagenomic insights into the microbial community of a thermal spring. Microbiome 7(8). https://doi.org/10.1186/s40168-019-0625-6

Peng X, Chen S, Xu H (2013) Formation of biogenic sheath-like Fe oxyhydroxides in a near-neutral pH hot spring: implications for the origin of microfossils in high-temperature, Fe-rich environments. Journal of Geophysical Research: Biogeosciences 118:1397–1413. https://doi.org/10.1002/jgrg.20119

Pierson BK, Castenholz RW (1974) A phototrophic gliding filamentous bacterium of hot springs, Chloroflexus aurantiacus, gen. and sp. nov. Archives of Microbiology 100:5–24. https://doi.org/10.1007/bf00446302

Pierson BK, Parenteau MN (2000) Phototrophs in high iron microbial mats: microstructure of mats in iron-depositing hot springs. FEMS Microbiol Ecol 32:181–196. https://doi.org/10.1111/j.1574-6941.2000.tb00711.x

Pierson BK, Parenteau MN, Griffin BM (1999) Phototrophs in high-iron-concentration microbial mats: physiological ecology of phototrophs in an iron-depositing hot spring. Appl Environ Microbiol 65:5474–5483. https://doi.org/10.1128/aem.65.12.5474-5483.1999

Porowski A, Porowska D, Halas S (2019) Identification of sulfate sources and biogeochemical processes in an aquifer affected by peatland: insights from monitoring the isotopic composition of groundwater sulfate in Kampinos National Park, Poland. Water. https://doi.org/10.3390/w11071388

Prieto-Barajas CM, Valencia-Cantero E, Santoyo G (2018) Microbial mat ecosystems: Structure types, functional diversity, and biotechnological application. Electronic Journal of Biotechnology 31:48–56. https://doi.org/10.1016/j.ejbt.2017.11.001

Pruesse E, Quast C, Knittel K et al (2007) SILVA: a comprehensive online resource for quality checked and aligned ribosomal RNA sequence data compatible with ARB. Nucleic Acids Res 35:7188–7196. https://doi.org/10.1093/nar/gkm864

Pruesse E, Peplies J, Glöckner FO (2012) SINA: accurate high throughput multiple sequence alignment of ribosomal RNA genes. Bioinformatics 28:1823–1829. https://doi.org/10.1093/bioinformatics/bts252

Reigstad LJ, Richter A, Daims H, Urich T, Schwark L, Schleper C (2008) Nitrification in terrestrial hot springs of Iceland and Kamchatka. FEMS Microbiol Ecol 64:167–174. https://doi.org/10.1111/j.1574-6941.2008.00466.x

Rozanov AS, Bryanskaya AV, Ivanisenko TV, Malup TK, Peltek SE (2017) Biodiversity of the microbial mat of the Garga hot spring. BMC Evolutionary Biology 17(2). https://doi.org/10.1186/s12862-017-1106-9

Sahoo RK, Subudhi E, Kumar M (2015) Investigation of bacterial diversity of hot springs of Odisha, India. Genomics Data 6:188–190. https://doi.org/10.1016/j.gdata.2015.09.018

Saxena R, Dhakan DB, Mittal P et al (2017) Metagenomic analysis of hot springs in central India reveals hydrocarbon degrading thermophiles and pathways essential for survival in extreme environments. Front Microbiol 7(2123). https://doi.org/10.3389/fmicb.2016.02123

Schloss PD, Westcott SL, Ryabin T et al (2009) Introducing mothur: open-source, platform-independent, community-supported software for describing and comparing microbial communities. Appl Environ Microbiol 75:7537–7541. https://doi.org/10.1128/AEM.01541-09

Schulz C, Schütte K, Koch N et al (2018) The active bacterial assemblages of the upper GI tract in individuals with and without Helicobacter infection. Gut 67:216–225. https://doi.org/10.1136/gutjnl-2016-312904

Sharma M, Polizzotto M, Anbar AD (2001) Iron isotopes in hot springs along the Juan de Fuca Ridge. Earth and Planetary Science Letters 194:39–51. https://doi.org/10.1016/s0012-821x(01)00538-6

Spanevello MD, Patel BKC (2004) The phylogenetic diversity of Thermus and Meiothermus from microbial mats of an Australian subsurface aquifer runoff channel. FEMS Microbiol Ecol 50:63–73. https://doi.org/10.1016/j.femsec.2004.05.008

Spietz RL, Williams CM, Rocap G, Horner-Devine MC (2015) A dissolved oxygen threshold for shifts in bacterial community structure in a seasonally hypoxic estuary. PLoS ONE 10(8): e0135731. https://doi.org/10.1371/journal.pone.0135731

Stieglmeier M, Klingl A, Alves RJE (2014) Nitrososphaera viennensis gen. nov., sp. nov., an aerobic and mesophilic, ammonia-oxidizing archaeon from soil and a member of the archaeal phylum Thaumarchaeota. Int J Syst Evol 64:2738–2752. https://doi.org/10.1099/ijs.0.063172-0

Takai K, Kobayashi H, Nealson KH, Horikoshi K (2003) Sulfurihydrogenibium subterraneum gen. nov., sp. nov., from a subsurface hot aquifer. Int J Syst Evol 53:823–827. https://doi.org/10.1099/ijs.0.02506-0

Tank M, Thiel V, Ward DM, Bryant DA (2017) A panoply of phototrophs: an overview of the thermophilic chlorophototrophs of the microbial mats of alkaline siliceous hot springs in Yellowstone National Park, WY, USA. In: Hallenbeck P. (ed) Modern topics in the phototrophic prokaryotes. Springer, Cham, pp 87–137

Thiel V, Hügler M, Ward DM, Bryant DA (2017) The dark side of the mushroom spring microbial mat: life in the shadow of chlorophototrophs. II. Metabolic functions of abundant community members predicted from metagenomic analyses. Front Microbiol 8(943). https://doi.org/10.3389/fmicb.2017.00943

Thiel V, Wood JM, Olsen MT et al (2016) Front Microbiol 7(919). https://doi.org/10.3389/fmicb.2016.00919

Uribe‐Lorío L, Brenes-Guillén L, Hernández‐Ascencio W et al (2019) The influence of temperature and pH on bacterial community composition of microbial mats in hot springs from Costa Rica. MicrobiologyOpen 2019(e893). https://doi.org/10.1002/mbo3.893

Valverde A, Tuffin M, Cowan DA (2012) Biogeography of bacterial communities in hot springs: a focus on the actinobacteria. Extremophiles 16:669–679. https://doi.org/10.1007/s00792-012-0465-9

Ward DM, Ferris MJ, Nold SC, Bateson MM (1998) A natural view of microbial biodiversity within hot spring cyanobacterial mat communities. Microbiology and Molecular Biology Reviews 62:1353–1370. https://doi.org/10.1128/mmbr.62.4.1353-1370.1998

Wang S, Hou W, Dong H et al (2013) Control of temperature on microbial community structure in hot springs of the Tibetan Plateau. PLoS ONE 8(5): e62901. https://doi.org/10.1371/journal.pone.0062901

Ward DM, Castenholz RW (2000) Cyanobacteria in geothermal habitats. In: Whitton BA, Potts M (eds) The ecology of cyanobacteria. Springer, Dordrecht, pp 37–59

Ward LM, Hemp J, Shih PM, McGlynn SE, Fischer WW (2018) Evolution of phototrophy in the Chloroflexi phylum driven by horizontal gene transfer. Frontiers in Microbiology 9(260). https://doi.org/10.3389/fmicb.2018.00260

Wu D, Raymond J, Wu M et al (2009) Complete genome sequence of the aerobic CO-oxidizing thermophile Thermomicrobium roseum. PLoS ONE 4(1): e4207. https://doi.org/10.1371/journal.pone.0004207

Xu Y, Schoonen MAA, Nordstrom DK, Cunningham KM, Ball JW (1998) Sulfur geochemistry of hydrothermal waters in Yellowstone National Park: I. the origin of thiosulfate in hot spring waters. Geochimica et Cosmochimica Acta 62:3729–3743. https://doi.org/10.1016/s0016-7037(98)00269-5

Xu Y, Schoonen MAA, Nordstrom DK, Cunningham KM, Ball JW (2000) Sulfur geochemistry of hydrothermal waters in Yellowstone National Park, Wyoming, USA. II. Formation and decomposition of thiosulfate and polythionate in Cinder Pool. J Volcanol Geotherm Res 97:407–423. https://doi.org/10.1016/s0377-0273(99)00173-0

Yarza P, Yilmaz P, Pruesse E et al (2014) Uniting the classification of cultured and uncultured bacteria and archaea using 16S rRNA gene sequences. Nat Rev Microbiol 12:635–645. https://doi.org/10.1038/nrmicro3330

Yee N, Phoenix VR, Konhauser KO, Benning LG, Ferris FG (2003) The effect of cyanobacteria on silica precipitation at neutral pH: implications for bacterial silicification in geothermal hot springs. Chemical Geology 199:83–90. https://doi.org/10.1016/s0009-2541(03)00120-7

Yilmaz P, Parfrey LW, Yarza P et al (2014) The SILVA and “All-species living rree project (LTP)” taxonomic frameworks. Nucleic Acids Res 42:D643–D648. https://doi.org/10.1093/nar/gkt1209

