## Supplementary information for "Rapid shift in microbial community structure in a neutral hydrothermal hot spring from Costa Rica"

Diego Rojas-Gätjens^1^, Alejandro Arce-Rodríguez^2,3,α^ , Fernando Puente-Sánchez^4^, , Roberto Avendaño^1^, Eduardo Libby^5^, Geraldine Conejo-Barboza^5^, Raul Mora-Amador^6^, Keilor Rojas^7^, Dietmar H. Pieper^3^ & Max Chavarría^1,5,8*^

^1^Centro Nacional de Innovaciones Biotecnológicas (CENIBiot), CeNAT-CONARE, 1174-1200, San José, Costa Rica ^2^Institute of Microbiology, Technical University of Braunschweig, D-38106, Braunschweig, Germany ^3^Microbial Interactions and Processes Research Group, Helmholtz Centre for Infection Research, 38124, Braunschweig, Germany ^4^Systems Biology Program, Centro Nacional de Biotecnología (CNB-CSIC), C/Darwin 3, 28049 Madrid, Spain ^5^Escuela de Química, Universidad de Costa Rica, 11501-2060, San José, Costa Rica. ^6^Escuela de Geología, Universidad de Costa Rica, 11501-2060, San José, Costa Rica ^7^Escuela de Biología, Universidad de Costa Rica, 11501-2060, San José, Costa Rica ^8^Centro de Investigaciones en Productos Naturales (CIPRONA), Universidad de Costa Rica, 11501-2060, San José, Costa Rica.

Keywords: Hot Spring, Costa Rica, Bajo las Peñas, Microbial Communities, Microbial mat, Ferryhidrite, Cross-feeding

^α^ Current affiliation:

Department of Molecular Bacteriology

Helmholtz Centre for Infection Research

38124, Braunschweig, Germany

^*^Correspondence to: Max Chavarría

Escuela de Química & Centro de Investigaciones en Productos Naturales (CIPRONA)

Universidad de Costa Rica

Sede Central, San Pedro de Montes de Oca

San José, 11501-2060, Costa Rica

Phone (+506) 2511 8520.  Fax (+506) 2253 5020

ORCID: <https://orcid.org/0000-0001-5901-3576>

**LEGENDS OF TABLES**

**Table S1. DNA sequence and phylogenetic assignment of the most abundant phylotypes detected in Bajo las Peñas using Illumina-based amplicon deep-sequencing.**

**See Excel file.**

**LEGENDS OF FIGURES**

**Fig. S1 Powder X-ray diffraction pattern for sediment samples of Bajo Las Peñas sites S1 and S2.** The red bars correspond to 6-line ferrihydrite as reported by Drits et al. (1993). X-ray diffractograms were acquired with a diffractometer (Bruker Powder D8 Advance), radiation source (Cu Kα1- Kα2) in a Bragg Bentano configuration, and a linear Lynx-eye detector.

**Fig. S2 Diversity measures of the samples in Bajo las Peñas.** The diversity measures (Observed Richness, Shannon and Simpson) were calculated using phyloseq. Figure shows A) diversity measures of all samples grouped by sample point. B) diversity measures of all samples.
