## Supplementary figures and images for "Rapid shift in microbial community structure in a neutral hydrothermal hot spring from Costa Rica"

### Supp. Figure S1

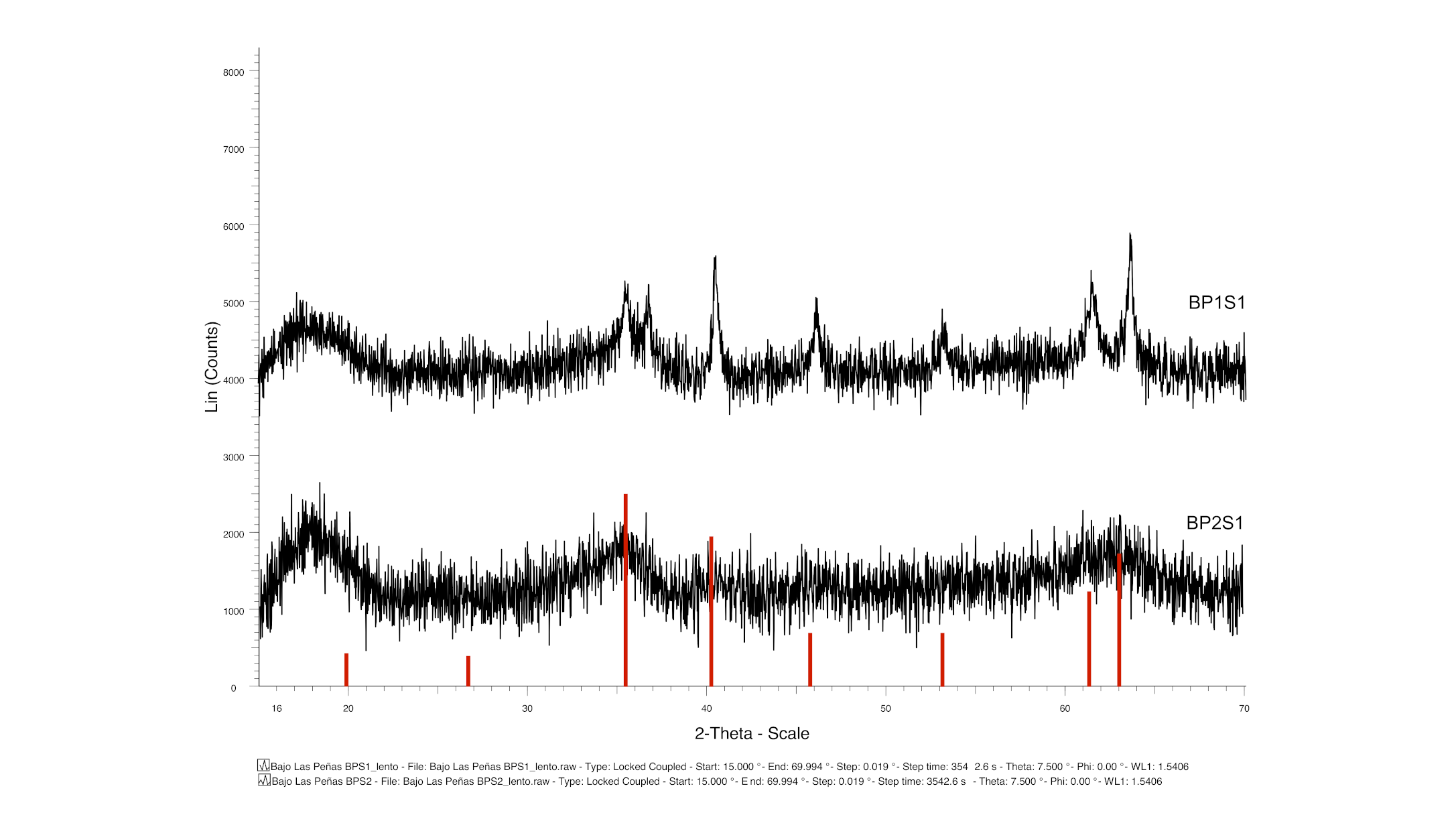

### Supp. Figure S2

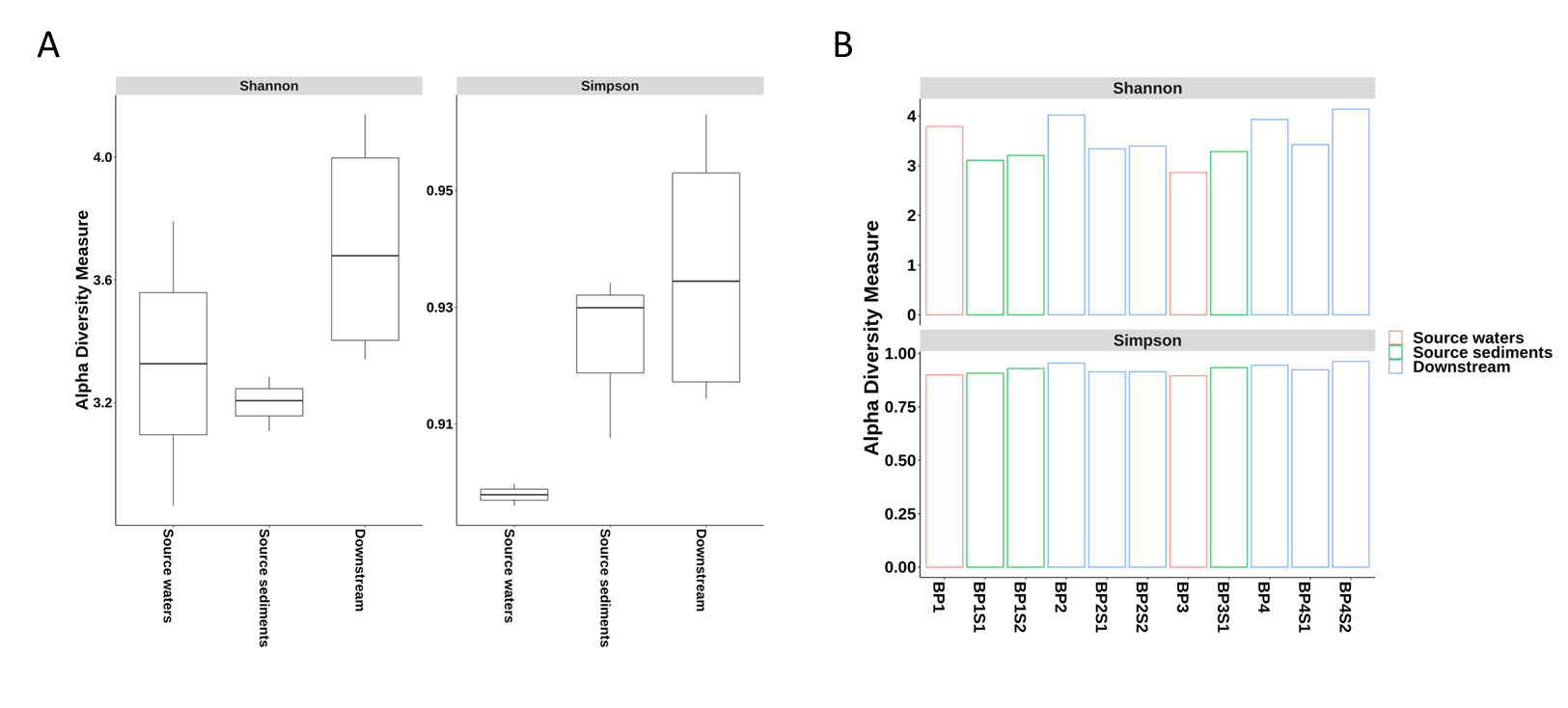
